# Gauge your phage: Benchmarking of bacteriophage identification tools in metagenomic sequencing data

**DOI:** 10.1101/2021.04.12.438782

**Authors:** Siu Fung Stanley Ho, Nicole Wheeler, Andrew D. Millard, Willem van Schaik

## Abstract

**Background:** The prediction of bacteriophage sequences in metagenomic datasets has become a topic of considerable interest, leading to the development of many novel bioinformatic tools. A comparative analysis of ten state-of-the-art phage identification tools was performed to inform their usage in microbiome research.

**Methods:** Artificial contigs generated from complete RefSeq genomes representing phages, plasmids, and chromosomes, and a previously sequenced mock community containing four phage species, were used to evaluate the precision, recall and F1-scores of the tools. We also generated a dataset of randomly shuffled sequences to quantify false positive calls. In addition, a set of previously simulated viromes was used to assess diversity bias in each tool’s output.

**Results:** VirSorter2 achieved the highest F1 score (0.92) in the RefSeq artificial contigs dataset, with several other tools also performing well. Kraken2 had the highest F1 score (0.86) in the mock community benchmark by a large margin (0.3 higher than DeepVirFinder in second place), mainly due to its high precision (0.96). Generally, k-mer based tools performed better than reference similarity tools and gene-based methods. Several tools, most notably PPR Meta, called a high number of false positives in the randomly shuffled sequences. When analysing the diversity of the genomes that each tool predicted from a virome set, most tools produced a viral genome set that had similar alpha and beta diversity patterns to the original population, with Seeker being a notable exception.

**Conclusions:** This study provides key metrics used to assess performance of phage detection tools, offers a framework for further comparison of additional viral discovery tools, and discusses optimal strategies for using these tools. We highlight that the choice of tool for identification of phages in metagenomic datasets, as well as their parameters, can bias the results and provide pointers for different use case scenarios. We have also made our benchmarking dataset available for download in order to facilitate future comparisons of phage identification tools.

## Introduction

Bacteriophages (phages) and archaeal viruses are globally ubiquitous, diverse and typically outnumber their prokaryotic hosts in most biomes [1]. Phages play a key role in microbial communities by: shaping and maintaining microbial ecology by fostering coevolutionary relationships [2]–[4]; biogeochemical cycling of essential nutrients [5]–[7]; and facilitating microbial evolution through horizontal gene transfer [8]–[10]. Despite the abundance and perceived influence phages have on all microbial ecosystems, they continue to be one of the least studied and understood members of complex microbiomes [11]. Phages are obligate parasites which require their bacterial host’s machinery to replicate, and spread via cell lysis. They can either be lytic or temperate, and while the former can only follow the lytic life cycle, temperate phages can either follow the lytic or lysogenic cycle [12]. During the lytic cycle, phages hijack host cell machinery to produce new viral particles. In the lysogenic cycle, phages can integrate their genomes into the bacterial host genome chromosome as linear DNA or as a self-replicating autonomous plasmid. In addition, a third life cycle called pseudolysogeny has been documented, in which either lytic phage infection is halted or there is no prophage formation [13].

Traditionally, phage identification and characterisation relied on isolation and culturing techniques, which are time consuming and often require significant expertise. It is also often impractical as many hosts, and their phages, cannot be cultured under laboratory conditions [14]. The arrival of high-throughput next generation sequencing has allowed metagenomic data from various environments to be generated routinely. Metagenomic sequencing allows direct identification and analysis of all genetic material in a sample, regardless of cultivability [15].

Metagenomic studies can opt to either sequence the whole community metagenome and then computationally isolate viral sequences, or physically separate the viral fraction before library preparation to produce a metavirome. The latter approach risks eliminating a large proportion of phages owing to their association with the cellular fraction. This occurs owing to phages being integrated into their hosts’ genome as prophages [16], attached to their hosts’ surface [17], or when they are in a pseudolysogenic state [18]–[20]. Purification methods may also remove certain types of phage, e.g. chloroform can inactivate lipid enveloped and/or filamentous phages [21], [22], increasing sampling bias. This process also results in low DNA yields, leading to some metavirome studies having to use multiple displacement amplification (MDA) to achieve sufficient quantities of DNA for library generation [11]. MDA has been shown to produce significant bias into virome composition [23], [24], by preferentially amplifying small circular ssDNA phage, such as those from the family *Microviridae* (Kim and Bae, 2011). Despite these drawbacks, the purification steps produce a metavirome with very little host contamination, although it is very difficult to produce a viral fraction that is devoid of any cellular material [25]. Metaviromes also have the advantage of being able to identify lower abundance phages at the same sequencing depth due to the approximately 100-fold larger bacterial genomes being excluded. Alternatively, whole community metagenomic sequencing can present insights into the host and viral fractions concurrently, allowing host-phage dynamics to be analysed. Integrated phages or prophages which have been found to be prevalent in some environments [26], can be identified since host genomes are also sequenced in this process. In this study, we focus on computationally extracting phage sequences from whole community metagenomes, as these generally make up a minority of the sequencing data compared to their bacterial hosts.

Many tools for identifying viral sequences from mixed metagenomic and virome assemblies have been developed in the last five years (Table 1). VirSorter [27] was one of the first of these, with previous tools focusing on prophage prediction (PhiSpy [28], Phage_Finder [29], PHAST/PHASTER [30], ProPhinder [31]) or virome analysis (MetaVir2) [32], VIROME [33]). VirSorter identifies phage sequences by detecting viral hallmark genes that have homology to reference databases, and by building probabilistic models based on different metrics (viral-like genes, PFAM genes, uncharacterised genes, short genes, and strand switching) which measure the confidence of each prediction. Since VirSorter’s release, other gene based homology based tools such as VIBRANT [35], and VirSorter2 [36] have been developed. VIBRANT uses a multilayer perceptron neural network based on protein annotation from Hidden Markov Model (HMM) hits to several databases to recover a diverse array of phages infecting bacteria and archaea including integrated prophages. In addition to this, it characterises auxiliary metabolic genes and pathways after identification. VirSorter2 builds on its predecessor by incorporating five distinct random forest classifiers for five different viral groups into one algorithm to improve the diversity of viruses that it can detect accurately. MetaPhinder, unlike the tools above, uses BLAST based homology hits to a custom database to calculate average nucleotide identity and the likelihood that a sequence is of viral origin.

**Table 1.** Overview of tools to identify and predict phage sequences in microbial ecosystems. Tools in italics were not included in this study as they were either not relevant to this study or technical difficulties were encountered during their use. MARVEL was excluded as it currently limited to detecting phages of the *Caudovirales* order.

| Software | Description | Reference |
| --- | --- | --- |
| DeepVirFinder | Predicts viral sequences via a k-mer based deep learning method using convolutional neural networks (CNN). Based on VirFinder. | [40] |
| *MARVEL* | Machine learning tool for predicting phage sequences in metagenomic bins. | [34] |
| MetaPhinder | Integrates BLAST hits to multiple phage genomes in a database to identify phage sequences in assembled contigs. | [45] |
| viralVerify<br>(metaviralSPAdes) | viralVerify is a module of metaviralSPAdes which classifies contigs with a Naïve Bayes classifier based on Hidden Markov Models protein hits. | [46] |
| *PhaMers* | Identifies phage sequences by a machine learning model based on k-mer frequencies. | [47] |
| PPR-Meta | Deep learning CNN approach to identify both phages and plasmids | [43] |
| Seeker | Deep learning framework that uses Long Short -Term Memory models (LSTM) which does not depend on sequence motifs. | [44] |
| VIBRANT | Deep learning neural network based on protein signatures which also highlights auxiliary metabolic genes and pathways. | [35] |
| *ViraMiner* | Extension of DeepVirFinder that is trained to identify any virus that may colonise human samples. | [48] |
| VirFinder | K-mer based machine learning method for identification of viral contigs. | [38] |
| *virMine* | Iterative pipeline that relies on the abundance of non-viral sequences in databases to strictly filter out unwanted contigs. Pipeline accepts both reads or assembled contigs. | [49] |
| *VirMiner* | Web-based pipeline that handles genome assembly, functional annotation using a variety of databases and identification of phage contigs via a random forest algorithm | [50] |
| *VirNet* | Deep learning neural network using an attentional neural model trained on nucleotide viral fragments. | [51] |
| *VIROME* | Web-based pipeline that classifies viral sequences based on homology to databases and functional annotates them. No local version. | [33] |
| VirSorter | Uses referenced-based and reference-free approaches in unison relying on probabilistic similarity models and referenced based protein homology searches to increase novel virus detection. | [27] |
| VirSorter2 | Builds on VirSorter by applying machine learning to evaluate “viralness” using genomic features. Works with a wider variety of viral groups than its predecessor. | [36] |
| *VirusSeeker* | Made up of two BLAST-based pipelines – Virome and Discovery. Virome aligns reads to a curated database to identify viral sequences and compute their abundance in the sample. Discovery focuses on contig-based analysis to aid novel virus discovery. | [52] |

VirFinder was the first machine learning, viral identification tool to utilise k-mer signatures [38]. VirFinder was shown to have considerably better rates of recovery of viral sequences than VirSorter, especially on shorter sequences (<5 kb), but had issues of variable performance in different environments, perhaps due to biases introduced by the reference data used for training the machine learning model [39]. DeepVirFinder [40] improves on VirFinder by applying a convolutional neural network that was trained on an enlarged dataset containing viral sequences from environmental metavirome sequencing data. DeepVirFinder boasts increased viral identification at all contig lengths over its predecessor VirFinder, whilst mitigating the latter’s bias towards phages that are easily cultivable in the laboratory. Kraken2 is a k-mer based metagenomic taxonomic classifier [41] that can be used for viral detection [42]. It queries k-mers to a database which associates it to their lowest common ancestor taxa which is then used to assign the taxonomic label. PPR Meta uses three convolutional neural network to identify if a sequence is of phage, plasmid, or chromosomal origin [43]. Sequence features are extracted by the network directly, instead of using pre-selected features such as k-mer signatures or genes. The three networks are also trained on three groups of different sequence lengths to improve its performance on shorter fragments, which some gene-based tools struggle with due to the low number of full-length genes available for analysis. Seeker also uses a neural network, in this case a Long Short-Term Memory (LSTM) model, which is not based on pre-selected features [44]. MetaviralSPAdes [46] uses an entirely different approach by leveraging variations in depth between viral and bacterial chromosomes in assembly graphs. The tool is split into three separate modules: a specialised assembler based on metaSPAdes (viralAssembly); a viral identification module that classifies contigs as viral/bacterial/uncertain using a Naive Bayesian classifier (viralVerify); and a module which calculates the similarity of a constructed viral contig to known viruses (viralComplete).

It is important to note that machine learning tools have the potential to identify novel species, which is especially important with the enormous diversity of phages that is theorised to still be unknown [53]. With the development of so many tools using a variety of approaches, a comprehensive comparison and benchmarking is needed to evaluate which tools are most applicable to researchers. The performance of each method can vary based on sample content, assembly method, sequence length, classification thresholds and other custom parameters. To address these issues, we have benchmarked ten metagenomic viral identification tools using both artificial contigs, mock communities and real samples.

## Results

### Benchmarking with RefSeq phage and non-viral artificial contigs

Ten commonly used tools for viral sequence identification in metagenomes were selected for evaluation: DeepVirFinder; Kraken2; MetaPhinder; PPR Meta; Seeker; VIBRANT; viralVerify; VirFinder; VirSorter; and VirSorter2. All of these tools can be run locally without reliance of a web server, accept metagenomic contigs as input, and have been published in the past decade.

We first evaluated all the programs on the same uniform datasets. All complete phage genomes deposited in RefSeq between 1 January 2020 and 12 August 2021 were downloaded, quality controlled, and fragmented to create a true positive set of artificial contigs. A negative set was constructed from all RefSeq bacterial and archaeal chromosomes and plasmids, submitted in the same time period. Multiple steps were taken to ensure these datasets did not include false positive or false negatives. First, all bacterial, archaeal, and phage RefSeq genomes deposited prior to 2020 and the training sets of each machine learning tool were used to dereplicate the datasets used in this study to remove any similar species that may cause overfitting of some tools. In addition, chromosome and plasmid sequences with ≥30% of their open reading frames having hits to the pVOG database were removed to exclude any remaining viral sequences. As the negative dataset was considerably larger than the positive dataset, we subsampled the negative set by 14.3-fold, resulting in 253 host chromosomes and 309 host plasmids (Table 2). This sampling rate was chosen to produce a phage:host ratio (∼1:19) that was similar to what is found in human gut microbiomes [53]. Finally, we removed chromosomes and plasmid sequences that contained integrated prophages using two state-of-the-art prophage detection tools, Phigaro [54] and PhageBoost [55], to prevent their erroneous identification as viral contigs. In total 2088 prophages were removed from the chromosome set and 91 from the plasmid set. The cleaned artificial contigs were then run on the benchmarking tools (Figure 1).

**Table 2.** Number of sequences at each stage of the RefSeq benchmarking workflow. Columns labelled with (n) contain the number of sequences at each step and columns with (bp) indicate the number of base pairs at each step. Steps are numbered as follows: 1. Sequences downloaded from RefSeq which were deposited between 1 January 2020 and 12 August 2021; 2. Sequences from (1) which were then dereplicated with RefSeq sequences deposited before 1^st^ January 2020, and training sets for DeepVirFinder, Seeker, VIBRANT, VirFinder, VirSorter2; 3. Host sequences (chromosome and plasmids) from (2) which were subsampled by a factor of 14.3; 4. Host sequences from (3) with prophage removal using Phigaro and PhageBoost. The number in parentheses indicates the number of prophages removed; 5. Host sequences from (4) with sequences that have ≥30% of their open reading frames having hits to the pVOG database removed; 6. All sequences from (5) randomly and uniformly fragmented to sizes between 1 and 15 kbp for use in the benchmarking study.

| Dataset | Chromosome (n) | Plasmid (n) | Phage (n) | Chromosome (bp) | Plasmid (bp) | Phage (bp) |
| --- | --- | --- | --- | --- | --- | --- |
| 1. Post-2020 | 7400 | 9960 | 1849 | 30,034,515,475 | 1,009,997,498 | 128,660,045 |
| 2. Post-2020 Dereplicated | 3546 | 4453 | 901 | 14,845,391,115 | 480,796,703 | 62,387,054 |
| 3. Subsampled | 253 | 309 | 901 | 1,011,740,231 | 28,782,822 | 62,387,054 |
| 4. Prophage Removed | 2307 (2,088) | 400 (91) | 901 | 965,959,872 | 27,323,488 | 62,387,054 |
| 5. pVOG removed | 2065 | 313 | 901 | 848,361,160 | 24,433,971 | 62,387,054 |
| 6. Fragmented artificial contigs | 104003 | 2754 | 6664 | 830,889,456 | 21,783,942 | 53,426,665 |

**Figure 1.**
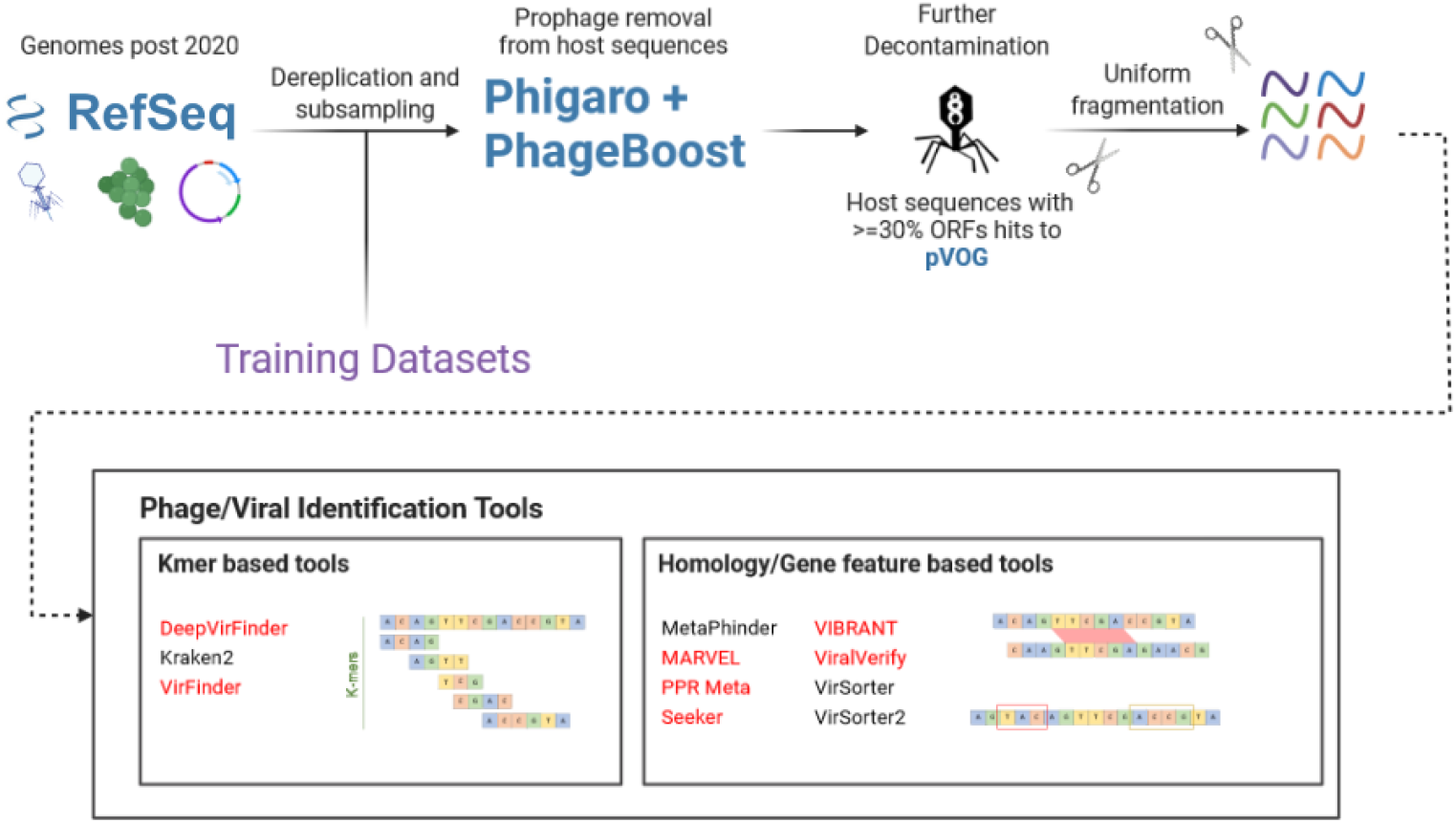
Overview of RefSeq benchmarking workflow. All bacterial and archaeal chromosomes and plasmids and phage genomes that were deposited in the RefSeq database between 1 January 2020 and 12 August 2021 inclusive were downloaded. The phage genomes were used to create a positive test set and the chromosomes and plasmids for a negative set. The sequences were dereplicated with the training sets for each machine/deep learning tool that was benchmarked (highlighted in red), as well as any RefSeq sequences deposited prior to 2020. The negative set was down sampled to produce a positive:negative ratio of approximately 1:19 to replicate a typical gut microbiome. Prophages were identified and removed with Phigaro and PhageBoost. Any host sequences with greater than 30% of open read frames having hits to the Prokaryotic Virus Orthologous Groups Database were then removed. All sequences were then uniformly fragmented into artificial contigs with lengths between 1 and 15 kbp. All identification tools were then run on the artificial contig sets.

All evaluated programs, except Kraken2, produce thresholds or confidence ranges for viral identification. For tools (DeepVirFinder, MetaPhinder, PPR Meta, Seeker, VirFinder, and VirSorter2) that assign a continuous threshold (score, identity, or probability), a F1 curve was plotted, and an optimal threshold was determined (Supplementary Figure 1). For VirSorter and viralVerify, the categories that returned the highest F1 score were used (Supplementary Figure 2). In most tools there was a trade-off between precision and recall. This is likely due to relaxed thresholds allowing for more viral and non-viral sequences to be detected, increasing recall, and decreasing precision simultaneously. For VIBRANT and VirSorter, the positive dataset was additionally run in virome mode and virome decontamination mode respectively, as this improves viral recovery in samples composed mainly of viral sequences by adjusting the tools sensitivity [27], [35]. The tools we benchmarked on this dataset had highly variable performance in terms of F1-score (0.44 – 0.93), precision (0.47 – 1.00), and recall (0.46 – 0.96) (Figure 2). VirSorter2 and PPR Meta achieved the highest F1-scores of 0.93 and 0.92 respectively. VirSorter2 achieved this with high precision (0.92) and well as high recall (0.93), whereas PPR Meta had a slightly lower precision at 0.88 but higher recall at 0.96. The majority of the remaining tools performed well, with six tools (DeepVirFinder, Kraken2, MetaPhinder, VIBRANT, VirFinder, and VirSorter) having F1-scores of over 0.83. Kraken2 had a precision score of almost 1, with only 2 chromosomal fragments and no plasmid fragments flagged as viral, whilst still correctly identifying over 5000 phage fragments. viralVerify had a low precision score of 0.55 whilst its recall score of 0.88 was comparable to the other tools. Seeker had poor performance in both precision (0.48) and recall (0.41) compared to the other tools. Generally, k-mer based tools performed better than reference similarity/gene-based tools, although the sample sizes of the investigated tools is too small to draw statistically significant conclusions. Across our benchmark, only 0.06% (4/6665) of phage contigs from our positive set were not detected by any tool, with 89.7% being identified by over half the tools, and 11.6% (775/6665) found by all 10 tools (Supplementary Figure 3).

**Figure 2:**
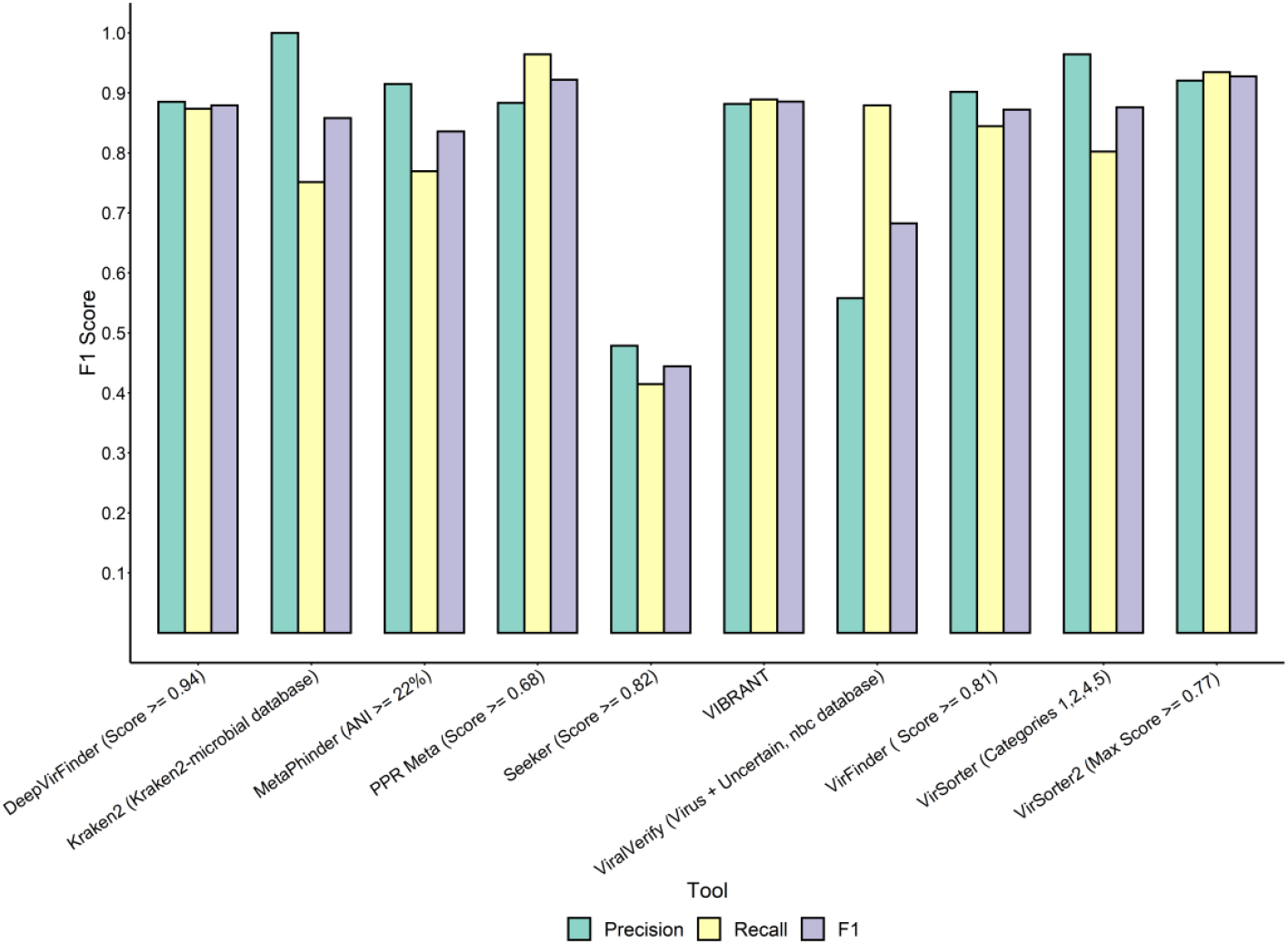
Comparison of viral identification tools on artificial RefSeq contigs. Contigs were generated by randomly fragmenting complete bacterial/archaeal/phage genomes and plasmids deposited in the NCBI Reference Sequence Database (RefSeq) between 1 January 2018 and 2 July 2020, to a uniform distribution. Each tool was then separately run on the true positive (phage genome fragments) and negative (bacterial/archaeal chromosome and plasmid fragments) datasets. For tools which score/probability threshold or categories could be manually adjusted, values/categories were selected based on optimal F1-scores.

### Benchmarking tools with randomly shuffled sequences

To serve as a further negative control, the positive RefSeq benchmark contigs were randomly shuffled at a nucleotide level to produce sequences that should not be identified as viral by any of the tools. Of the tools tested, four identified zero shuffled contigs (MetaPhinder, viralVerify, VirSorter, and VirSorter2), Kraken2 classified three contigs, DeepVirFinder and VirFinder detecting 742 and 1070 respectively, with the rest of the tools identifying over 2500 shuffled contigs as viral, including PPR Meta which incorrectly classifying 99.2% (6608/6664) of all the shuffled contigs as viral (Table 3).

**Table 3.**
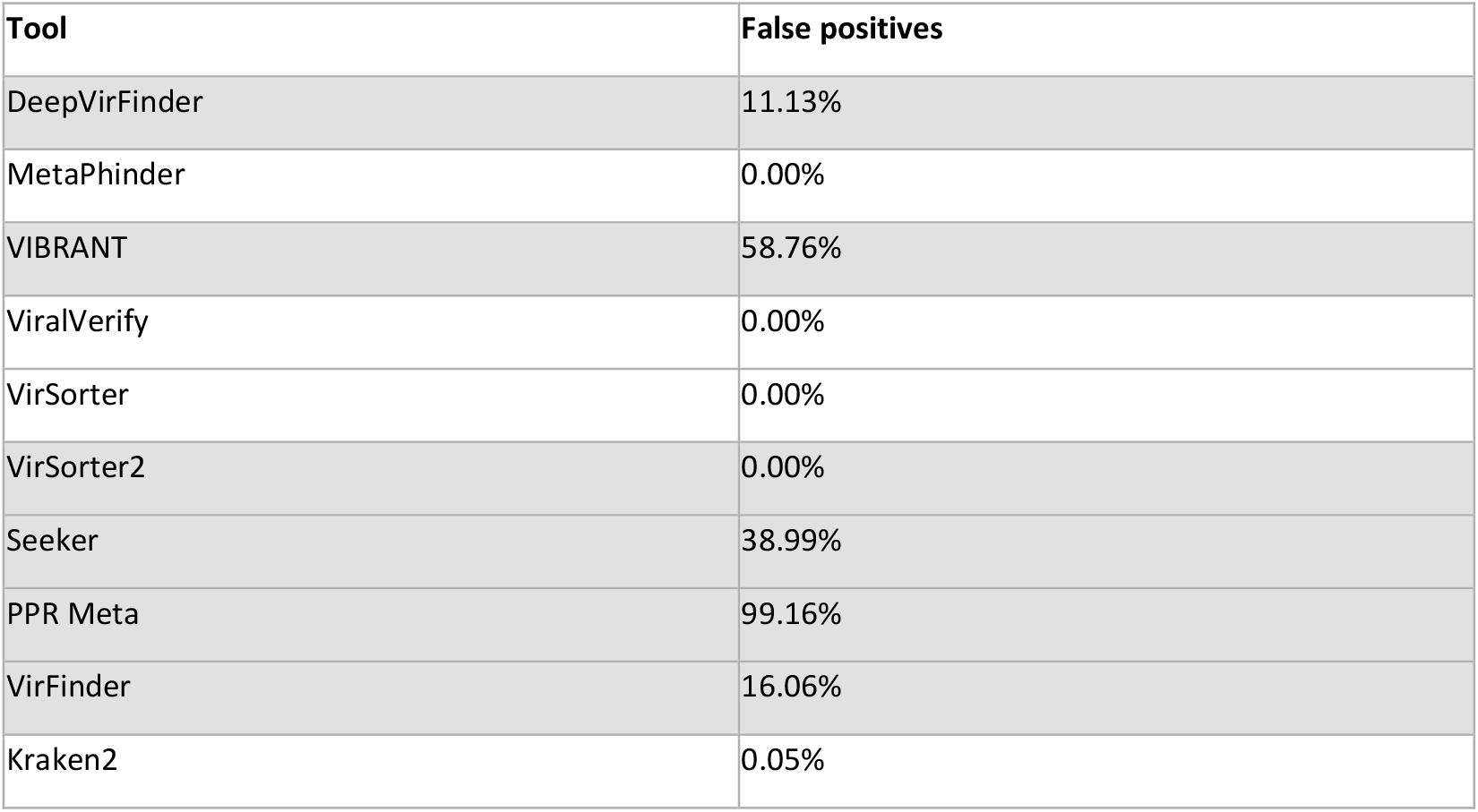
Performance of tools on randomly shuffled artificial phage contigs. Artificial RefSeq phage contigs generated in the previous benchmark were randomly shuffled whilst preserving the dinucleotide distribution using esl-shuffle from the HMMER3 suite. Phage detection tools were then run on the shuffled contigs, and any positive hits were recorded. Tools shaded in grey indicate methods utilising machine/deep learning models.

| Tool | False positives |
| --- | --- |
| DeepVirFinder | 11.13% |
| MetaPhinder | 0.00% |
| VIBRANT | 58.76% |
| ViralVerify | 0.00% |
| VirSorter | 0.00% |
| VirSorter2 | 0.00% |
| Seeker | 38.99% |
| PPR Meta | 99.16% |
| VirFinder | 16.06% |
| Kraken2 | 0.05% |

### Benchmarking tools with mock community shogun metagenomes

We next sought to compare these tools on real community shotgun metagenomic contigs. Thus, we obtained sequencing data of an uneven mock community created by Kleiner *et al*. [56], containing 32 species from across the tree of life, including five bacteriophages, at a large range of cell abundances (0.25%-21.25%; Supplementary Table 1). This allowed us to assess the performance of our tools on real data whilst retaining knowledge of the ground truth (sample composition) and determine each tool’s detection limit on low abundance species. The optimised parameters found in the RefSeq benchmark were used for each tool, with the exception of viralVerify and VirSorter. These tools have categorical thresholds which drastically change the profile of identified viral contigs, and the F1-scores between the thresholds were very close, so the parameters were further analysed in this benchmark. In general, the tools’ F1-scores were considerably lower on this dataset than on the RefSeq artificial contigs, with F1-scores dropping by an average of 40.6%, compared to the RefSeq benchmark (Figure 3). Kraken2 outperformed all other tools with a F1-score of 0.86, 0.3 higher than DeepVirFinder in second place. This is due to its high precision of 0.99 whilst identifying 76% of the phage contigs. DeepVirFinder had a high recall rate (0.80), but unlike in the RefSeq benchmark, had a lower precision of 0.42. Several other tools had similar results, with PPR Meta, MetaPhinder, and VirFinder all achieving high precisions of 0.91, 0.98, and 0.83 respectively, but having comparatively low precision scores (0.24, 0.28, and 0.35 respectively). VirSorter (categories 1, 2, 4, and 5) and Seeker had comparable performances to the tools above, with them both having F1-scores of 0.44. Seeker, along with Kraken2, were the only tools to attain similar F1-scores in this benchmark to the RefSeq benchmark. VirSorter2, which performed best in the RefSeq benchmark, had a lower F1-score (0.36) than its predecessor mainly due to its high number of false positive hits. VIBRANT, which also performed well in the previous benchmark, again had poor precision and middling recall resulting in a F1-score of 0.32. viralVerify (NBC database, virus only) had both low precision (0.24) and recall (0.22) resulting in the lowest F1-score (0.22) in this test. K-mer tools on average had a higher F1-score than the reference similarity/gene based tools but this difference was not statistically significant due to the small sample size of the number of tools in this study (*p = 0*.*094*, one-tailed Welch’s t-test).

**Figure 3:**
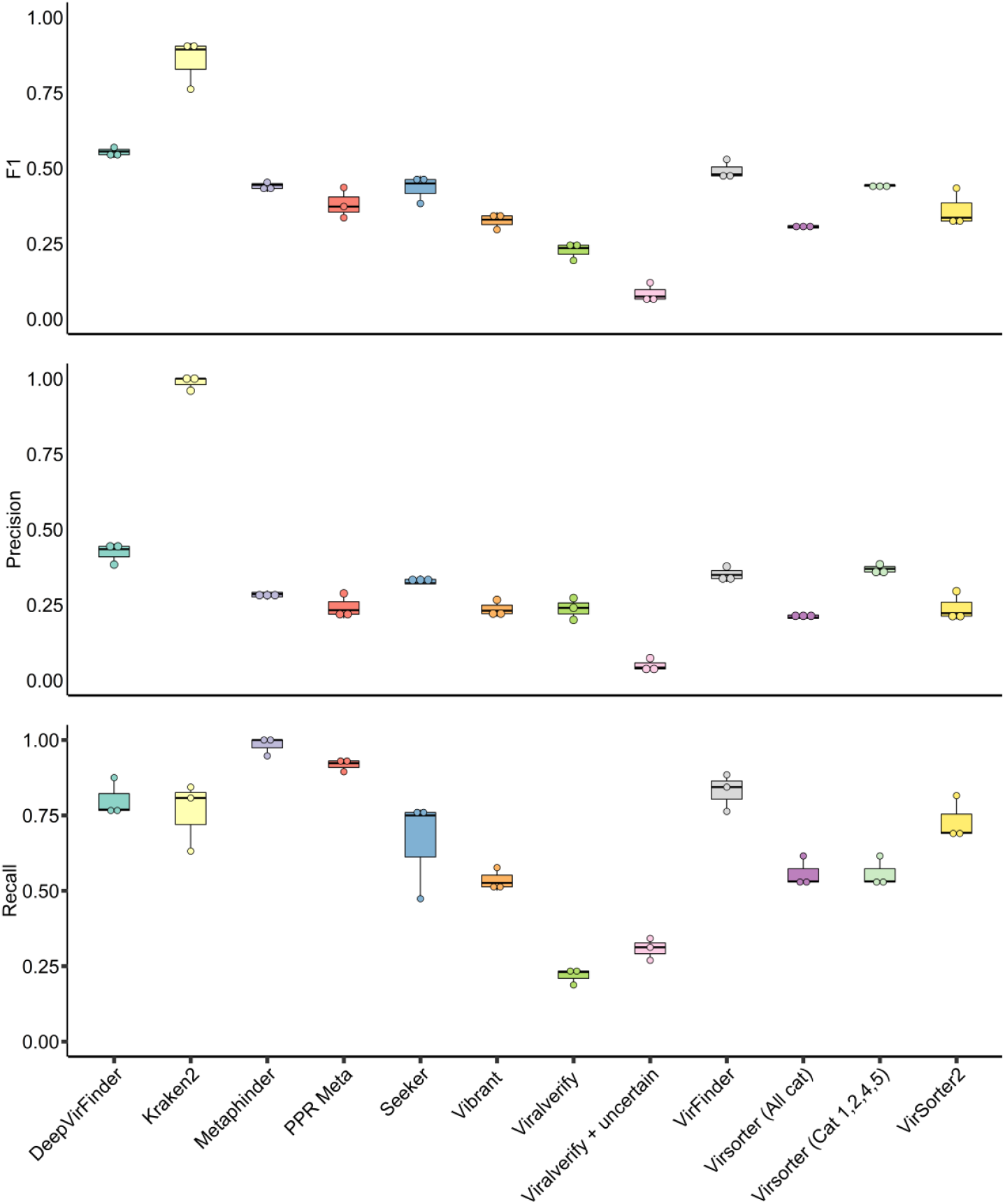
Comparison of viral identification tools on uneven mock community samples. Mock community reads were retrieved from a previous study [56]. and assembled with metaSPAdes before running each identification tool using optimal thresholds based on previous benchmarks with the exception of viralVerify and VirSorter. F1-score, Precision, and Recall metrics are displayed as separate panels. Each sample is plotted as a single point for each tool, with a boxplot indicating the interquartile ranges, extremes and mean of all three samples.

Out of the four DNA phage species found in the assemblies (Phage F2 was not sequenced due to being a ssRNA virus) only MetaPhinder was able to detect M13, whilst F0, the most abundant phage species, was detected by all benchmarked tools in all samples.

MetaPhinder identified contigs belonging to all four phage species in two samples and three in the other sample. PPR Meta, VIBRANT, VirSorter, and VirSorter2 were able to identify contigs belonging to three species in all three samples. viralVerify and VirFinder were able to identify the three phage species in two out of three samples, missing out on contigs belonging to phage ES18. DeepVirFinder and Kraken2 classified viral contigs belonging to three phage species in one out of three samples, and detected two species in the other samples. Seeker was only able to identify contigs belonging to the most abundant phage F0. No correlation was found between F1-score and the number of phage strains detected (*R*_*s*_ = -0.371, *p =* 0.29) but a positive, but not statistically significant, correlation, was observed between tools that identified more contigs of viral origin (true positives + false positives), and the number of phage strains identified (*R*_*s*_ = 0.604, *p* = 0.06).

### Impact of tool prediction on diversity metric estimation

To test the impact of these tools on diversity estimations, four simulated mock community metaviromes containing an average of 719 viral genomes were retrieved from [57]. Reads were mapped to contigs (>1 kb) that were identified as viral by each tool, and these mapped reads were then mapped to a set of population contigs to estimate their abundance in each sample. Original reads were also directly mapped to the population contigs as a control. Read counts were then normalised by their length and sequencing depth, which Roux *et al*. [57] found to be reliable normalisation method. Diversity estimation metrics were then calculated using the normalised population counts. All tools returned fewer genomes per sample compared to the initial population, although there was significant variation between tools. PPR Meta, MetaPhinder, and Kraken2 retrieved the greatest percentage of genomes with 86.8%, 89.1%, and 83.7% respectively (Figure 4A). All other tools were able to retrieve more than 50% of the genomes with the exception of Seeker and viralVerify, which were only able to recover 32.4% and 41.3%, respectively, of the population genomes. All Shannon’s alpha diversities calculated from the count matrices of each tool were within 10% of the default population with the exception of Seeker, whose *H* score was on average 27.0% lower (Figure 4B). Simpson alpha diversity indices showed similar performance, with all tools having a diversity score within 1% of the initial population, with the exceptions of DeepVirFinder and Seeker, who were 1.4% and 5.1% divergent, respectively (Figure 4C). PPR Meta was the only tool to estimate a comparatively higher alpha diversity than the default population. For beta diversity, pairwise Bray-Curtis dissimilarities within a sample were small between all tools except Seeker, whose Analysis of Similarity (ANOSIM) showed significant dissimilarity when compared to other tools (r = 0.3926, *p = 0*.*0019* with Benjamini–Hochberg correction for multiple comparisons at FDR = 0.05) (Figure 4D; Supplementary Table 2).

**Figure 4:**
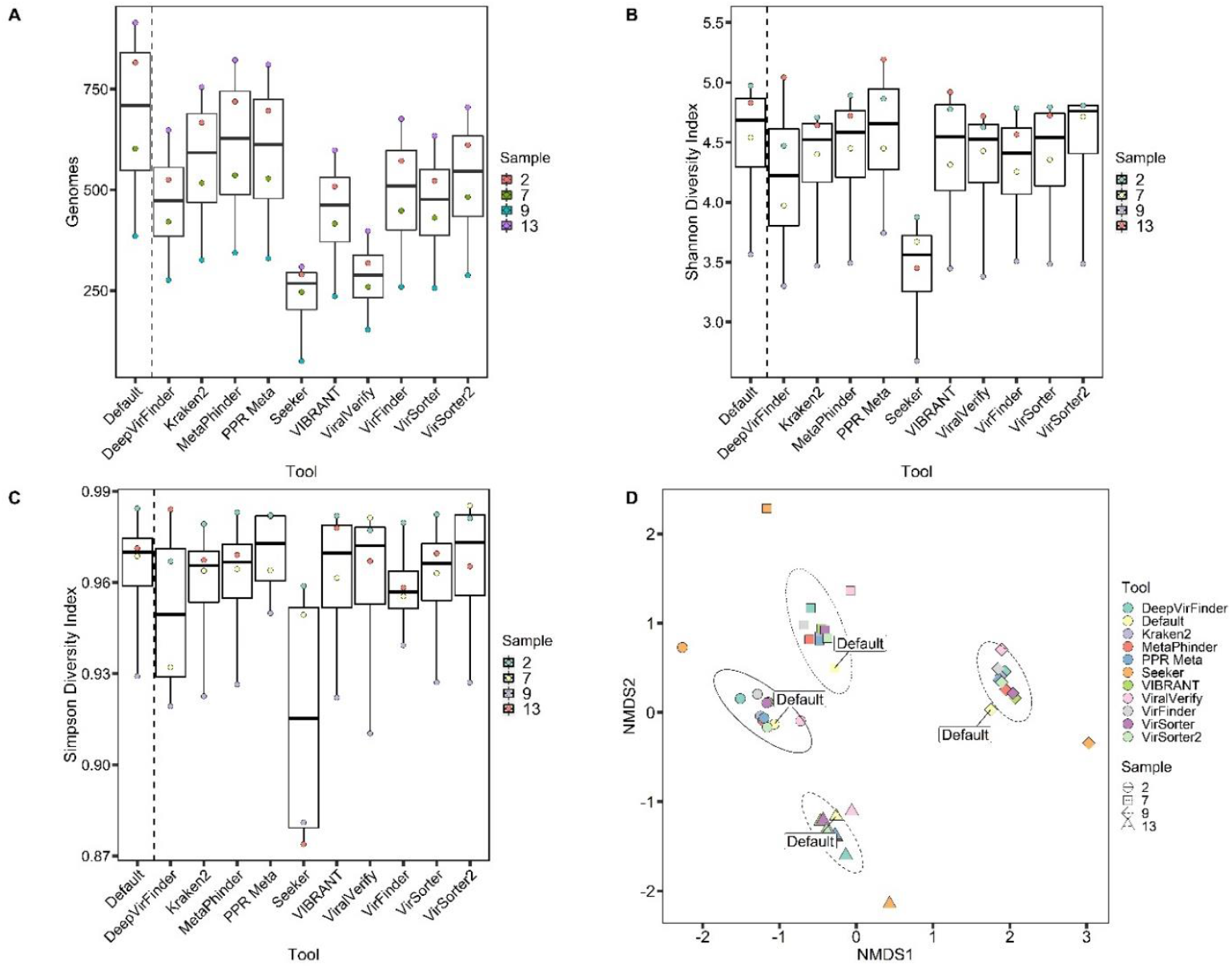
Estimation of diversity metrics of tool predicted virome populations. To assess the impact of each tool on population diversity, four simulated virome assemblies from Roux et al. were downloaded. Each program was then run to determine the subset of predicted viral contigs. Reads were mapped to these contig subsets and mapped reads were then subsequently mapped to a pool of population contigs. All diversity metrics were computed by the R package “vegan”. ‘Default’ in each plot indicates each sample’s original assembly. A. Number of genomes observed from read mapping to predicted viral contig populations for each tool. B. Comparison of estimated Shannon diversity indices from each tool’s virome subset. Estimations are based on read counts that were normalised by contig size and sequencing depth of the virome. C. Comparison of Simpson diversity indices from each tool’s virome subset. D. Non-metric multidimensional scaling (NMDS) ordination plot of Bray-Curtis dissimilarity of virome subsets predicted by each viral identification tool. Ellipses indicate the 95% confidence interval for each sample cluster’s centroid. Samples are represented by the same symbol and ellipse line type; tools are denoted by colour.

### Runtime and computational load of each tool

We also recorded the running times of each tool on the RefSeq positive dataset on a high-performance cluster (16 VCPU, 108 Gb RAM) (Figure 5). Kraken2 and VirFinder were the fastest tools finishing in under 5 minutes. DeepVirFinder, MetaPhinder, PPR Meta, and Seeker finished in under half an hour with viralVerify finishing just over that mark. VirSorter and VirSorter2 took the longest time to run on this dataset (2.9 hours and 3.8 hours to completion, respectively).

**Figure 5:**
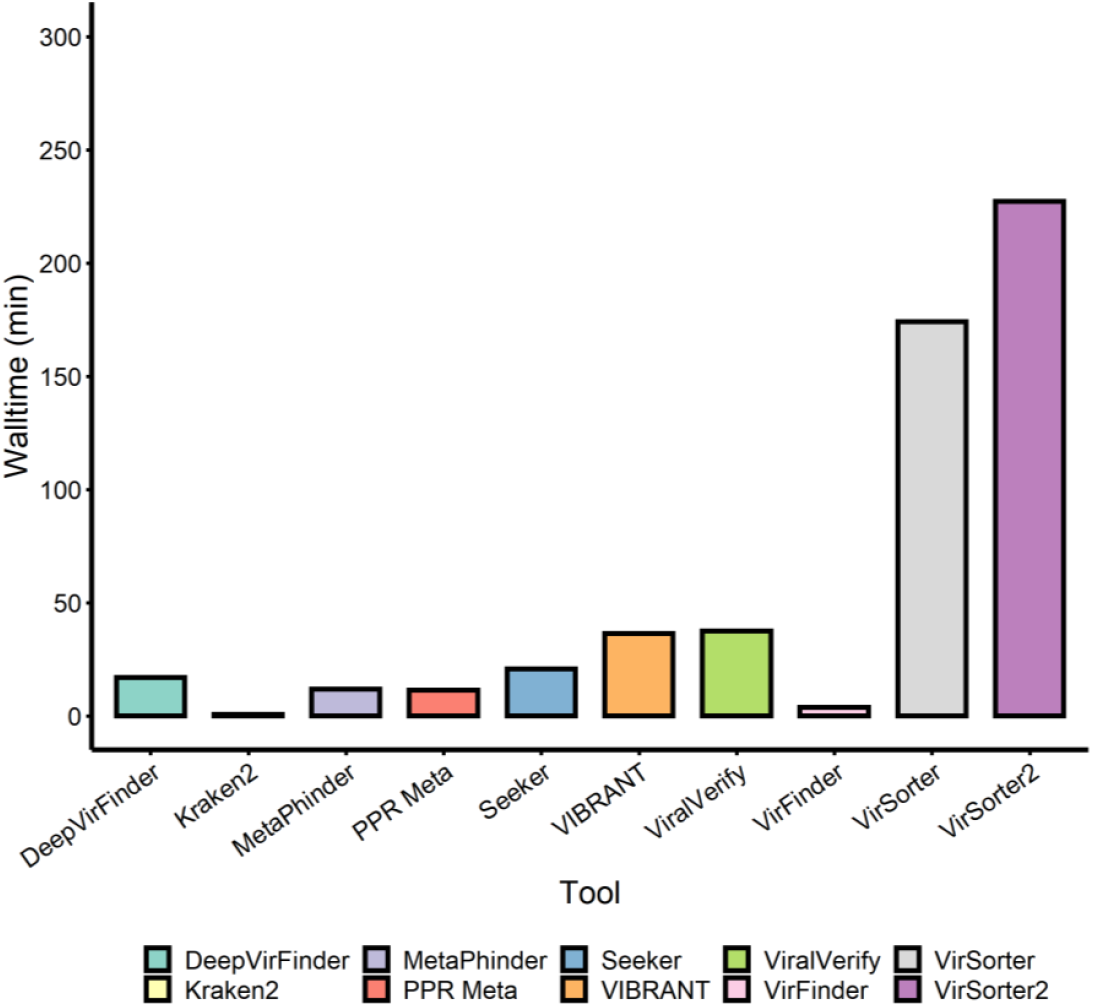
Comparison of tool runtimes on the positive RefSeq artificial contig set. Wall runtime of each tool on mock community samples was recorded on a 16 VCPU, 108GB RAM, Linux high performance cluster without GPU acceleration. Tools were run with 16 threads were possible. The RefSeq positive set contains approximately 53.4 million bp.

## Discussion

Bacteriophages are crucial members of microbial communities in nearly every ecosystem on Earth, responsible for controlling host population size as well as having wider impacts on community functions. Tools designed to recover viral sequences from mixed community metagenomic and virome samples are fundamental to studying the role of bacteriophages in the wider context of their environment. Advancements in this field have produced an extensive suite of viral identification tools that each claim to improve on the performance of similar tools. Selecting which tool among these is ideal for a dataset is thus not straightforward, especially as each novel tool typically only benchmarks against two or three other existing tools. Most tools developed for this purpose, especially those released in recent years have utilised machine/deep learning to classify sequences, whereas others rely on categorising sequences based on their similarity to sequences in databases. Both these approaches have potential to improve over time with newly discovered viral genomes being added to training datasets and databases.

Here we compare ten methods for identifying viral sequences from metagenomes across three datasets. We first benchmarked the tools on positive and negative datasets to evaluate their performance on an ideal set of contigs (size between 1 and 15 kb, without mis-assemblies), and determine approximate optimal thresholds. Most tools performed well here, detecting the majority of phage sequences, whilst keeping false positives low. PPR Meta and VirSorter2, which use two different machine learning methods, had the best performance across the tools. Generally, k-mer tools outperformed reference similarity and gene-based tools. Whilst the optimal thresholds that we determined may not necessarily be ideal for all other datasets, we believe they can be used as a basis for further usage of these tools as in each case they produced considerably better results than the default parameters. We therefore encourage researchers to apply these thresholds and parameters within the context of their prospective dataset.

A stark contrast between machine learning and more traditional tools can be seen when analysing their identification of randomly shuffled phage sequences. Of the six tools that utilise machine/deep learning methods, five identified a significant proportion of the sequences as viral, with only VirSorter2 as the only exception, probably due to its classifier being trained on a range of sequence and gene features. Three of the other four tools returned zero false positives and Kraken2 only returned three. This highlights that whilst these machine/deep learning algorithms have the capability to detect novel phages, their performance may be unpredictable when exposed to novel data with features that differ from those in our training sets.

When tested on real metagenomic data, most tools performed significantly worse than in the RefSeq benchmark, with the exceptions of Kraken2 and Seeker. Generally, k-mer tools had a smaller drop in F1-score from their RefSeq benchmark compared to reference similarity/gene-based tools. This is probably due to the comparatively shorter phage sequences that were assembled from the metagenomic reads, which provide less genetic context, thereby negatively impacting the algorithms of the reference similarity and gene-based tools. MetaPhinder was the only tool which was able to detect the one contig of phage M13 that was assembled in each sample. Unfortunately, this is likely due to MetaPhinder’s low precision in this benchmark resulting in many false positive calls. Most other tools were able to identify contigs belonging to the other three phages, with the exception of Seeker which could only identify the most abundant phage F0 in each sample. This suggests that when analysing metagenomic datasets where phage species are likely to be in low abundance, k-mer based tools such as Kraken2 and DeepVirFinder are the best choices, the former being the favoured option when precision is of particular importance, whilst the latter’s use is appropriate if the discovery of novel phages is of interest due to its deep learning algorithm.

We also gauged any potential biases and impact these tools may have on the diversity of its predicted viral population. Most tools performed well with alpha diversity indices within 10% of the default population with the exception of Seeker which returned a considerably lower value due to the very low number of viral population genomes Seeker originally predicted. Some tools such as PPR Meta predicted higher alpha diversity than default population. This is due to the tools missing some high abundance genomes from their predictions, resulting in a more even diversity distribution. When evaluating beta diversity, Seeker was the only tool that produced results that had significant dissimilarity from the other tools and did not cluster with the other programs, again as a result of the low proportion of genomes it recovered in this dataset. Hence beta diversity trends of the tools examined here, with the exception of Seeker, are accurate to the original population, even when only half the genomes are recovered.

Runtime and computational load are also important factors to examine, since these can become practical limitations if large samples take many hours or days to be analysed. Most tools were reasonably fast, although VirSorter, and its successor VirSorter2, took multiple hours to complete their runs. It is important to note that VIBRANT, VirSorter, and VirSorter2 annotate the identified viral genomes and predict prophages which comes at the expense of runtime, although these can be useful for some applications. Kraken2 was by far the fastest tool, taking less than a minute to run on our dataset. However, Kraken2 requires very high RAM use compared to the other benchmarked tools so it may not be feasible for researchers with limited computing power.

Although these benchmarks comprehensively compared the performance of state-of-the-art tools, there are a number of limitations with our study. First, whilst we use RefSeq genomes, and a mock metagenomic community to benchmark these tools, we do not address the tools’ ability to identify viral sequences belonging to different phage families. Secondly, we used the default database(s) or the original trained model(s) that was provided with each tool. Whilst providing each tool the same database, or dataset to be trained on, may have been a fairer comparison of the underlying algorithms, this was beyond the scope of our study. We note that most routine users are also unlikely to retrain these tools prior to their use. Thirdly, we did not assess the performance of combining multiple tools, which could provide meaning insights that would be missed when only one single tool is used, as in Marquet *et al*. [58] where the authors combined multiple tools into a single workflow. Fourth, many of the tools have additional functionalities, which we did not benchmark here but may nevertheless impact a researcher’s choice of tool such as: prophage prediction (VIBRANT, VirSorter, VirSorter2), plasmid prediction (PPR Meta); taxonomic identification (Kraken2); and functional annotation (VIBRANT). Finally, a few recently developed tools we found during our study were not included in our benchmarking either due to (1) requiring the use of its own web server and therefore not being scalable (VIROME, VirMiner), (2) lack of clear installation/running instructions (ViraMiner), or (3) errors when attempting to use the tool that we were unable to resolve (PhaMers, VirNet, VirMine).

## Conclusion

Our comparative analysis of ten currently available metagenomic virus/phage identification tools provides valuable metrics, and insights for other investigators to use and build on. Using mock communities and artifical datasets, precision, recall and biases of these tools could be calculated. By adjusting the filtering thresholds for viral identification for each tool and comparing F1 scores, we were able to optimise performance in every case. In the artificial RefSeq contig benchmark, most tools performed well, with PPR Meta and VirSorter2 having the highest F1-scores. In the mock uneven community dataset, tools generally performed worse with the exception of Kraken2, whose performance included almost perfect precision with above average recall scores. All tools except Seeker were able to produce a diversity profile with similar indices to the original virome population, and are therefore suitable for phage ecology studies. We suggest that of the currently available metagenomic phage identification tools, Kraken2 should be considered when researchers are trying to identify previously characterised phages. When novel phage detection is required Kraken2 should be used in combination with tools such as VirSorter2 and DeepVirFinder.

## Materials and Methods

### Benchmarking with RefSeq dataset

Complete bacterial and archaeal chromosome and plasmid sequences, and phage genomes deposited in RefSeq [59] since 1 January 2020 (inclusive) were downloaded on 12 August 2021 to construct a benchmarking set. Sequences in this set with ≥95% identity to the pre-2020 RefSeq sequences and training datasets for DeepVirFinder, PPR Meta, Seeker, VIBRANT, VirFinder, and VirSorter were removed with dedupe.sh [60] to reduce any potential overfitting. Chromosome and plasmid sequences were then randomly down sampled by a factor of 14.3, using reformat.sh (from BBtools suite) [60], to produce a host:phage ratio of approximately 19:1. Phigaro (v2.3.0, default settings) [54] and PhageBoost (v0.1.7, default settings) [55], were run in succession on the chromosomal and plasmid sequences to remove prophage sequences. Host sequences with ≥30% open reading frames (ORFs) with HMM hits to the pVOG database were removed as contamination. All sequences were then uniformly fragmented to between 1 kb and 15 kb, using a custom python script (available at https://github.com/sxh1136/Phage_tools), to create artificial contigs. Each viral prediction tool was then run on the three sets of contigs (chromosome, plasmid, and phage) with default settings except for VIBRANT and VirSorter where the phage-derived contig set was additionally run using their virome decontamination modes, due to their potentially improved performance in datasets consisting of mainly viral fragments. Commands used to run each tool and their version numbers can be found at https://github.com/sxh1136/Phage_tools/blob/master/manuscript_tools_script.md. RefSeq benchmarking datasets are available for download and use at https://figshare.com/articles/dataset/RefSeq_Datasets_for_benchmarking_-_Ho_et_al_/19739884.

For tools where score/probability thresholds can be manually adjusted (DeepVirFinder, MetaPhinder, PPR Meta, Seeker and VirFinder), F1 curves were plotted (100 data points) and optimal thresholds were determined by maximal F1 score. viralVerify and VirSorter, which have categorical thresholds for phage identification, the category sets with the highest F1-score was plotted; for VirSorter two category sets were compared, category 1,2,4,5 and all categories, as these are commonly used. viralVerify was also additionally run using both Pfam-A 33.0 and a provided database of virus/chromosome specific HMMs as these are listed on the tool’s GitHub usage guide.

Run time of each tool on this dataset, utilising 16 threads, was recorded using a Linux virtual machine provided by Cloud Infrastructure for Big Data Microbial Bioinformatics (CLIMB-BIG-DATA), with the following configuration: CPU: Intel® Xeon® Processor E3-12xx v2 (8 VCPU); GPU: Cirrus Logic GD 5446; Memory: 64GB Multi-Bit ECC.

### Benchmarking with randomly shuffled sequences

All artificial phage contigs created in the previous benchmark were randomly shuffled whilst preserving dinucleotide distribution using esl-shuffle from the HMMER3 suite (v3.3.2, -d) [61] Each identification tool was then run on the randomly shuffled sequences using the optimised thresholds that were determined in the RefSeq benchmark, and false positives were recorded.

### Benchmarking with mock community metagenomes

Three shotgun metagenomic sequencing replicates of an uneven mock community [56] was retrieved from the European Nucleotide Archive (BioProject PRJEB19901). These communities contain five phage strains: ES18 (H1), F0, F2, M13, and P22 (HT105). The quality of the data was checked using FASTQC (v0.11.8, default settings) [62] and overrepresented sequences were removed with Cutadapt (v2.10, --max-n 0) [63]. Cleaned paired-end reads were then assembled with MetaSPAdes (v3.14.1, default settings) [64] and contigs <1 kb were removed. Each tool was then run on the three sets of contigs using optimal parameters as determined previously with the exception of viralVerify and VirSorter where all categorical thresholds were re-evaluated. MetaQUAST (v5.0.2, default settings) [65] was used to map contigs to reference phage genomes and calculate coverage.

### Benchmarking with simulated mock virome communities

Four mock communities (samples 13, 2, 7, and 9) containing between 500 and 1000 viral genomes created by Roux *et al*. were selected for analysis [57]. These samples belonged to four different beta diversity groups and did not share any of their 50 most abundant viruses. Each simulation of 10 million paired-end reads were quality controlled with Trimmomatic [66] and assembled with MetaSPAdes by Roux *et al*. [57]. The contigs were then downloaded for benchmarking. As before, contigs with length <1 kbp were removed, and then inputted into each viral identification program. Positive viral contig sets for each tool were then extracted and reads were mapped to these with BBMap [67] with ambiguous mapped reads assigned to contigs at random (ambiguous=random), as in Roux *et al*. [57]. Primary mapped reads with pairs mapping to the same contig (options -F 0×2 0×904) were then extracted with SAMtools (v1.11) and mapped to a pool of non-redundant population contigs. This pool was created by clustering all four samples with nucmer (v3.1) [68], at ≥95% ANI (average nucleotide identity) across ≥80% of their lengths. Abundance matrices for each tool were calculated by normalising read counts by contig length and total library size to produce counts per million (CPM).These abundance matrices were then used to calculate Shannon, Simpson, and Bray-Curtis dissimilarity indices using the vegan package (v2.5.7) [69]. Non-metric multidimensional scaling (NMDS) and analysis of similarity (ANOSIM) were also computed with vegan. ANOSIM *p-*values were corrected with the Benjamini–Hochberg method [70]. Seed and permutations were set as 123 and 9999 respectively, where possible. All plots were generated with ggplot2 (v3.3.2) [71] and arranged with ggarrange from ggpubr (v0.4.0) [72].

## Abbreviations

ANOSIM: Analysis of Similarity
CNN: Convolutional Neural Networks
GPU: Graphics Processing Unit
kb: Kilobase pairs
MDA: Multiple Displacement Amplification
NCBI: National Center for Biotechnology Information
NMDS: Non-metric multidimensional scaling
RefSeq: NCBI Reference Sequence Database
ssDNA: Single Stranded DNA
TMM: Trimmed mean of M values
TPM: Transcripts Per Kilobase Million
VCPU: Virtual Central Processing Unit

## Availability of data and materials

RefSeq dataset construction and example commands used to run each tool are available at https://github.com/sxh1136/Phage_tools. RefSeq benchmarking datasets are available for download and use at https://figshare.com/articles/dataset/RefSeq_Datasets_for_benchmarking_-_Ho_et_al_/19739884

## Notes

https://github.com/sxh1136/Phage_tools

https://figshare.com/articles/dataset/RefSeq_Datasets_for_benchmarking_-_Ho_et_al_/19739884

### Competing Interest Statement

The authors have declared no competing interest.

### Summary of Updates

In this revised version, we have made the following changes: We improved the filtering of the databases we used for benchmarking to reduce contaminant sequences, particularly in the phage DNA database. We also removed sequences that overlapped with the training sets that were used for the different tools as that may be another factor that skew the outcome of the benchmarking analyses. An outline of this approach is now provided in Fig. 1. We included a true negative control dataset of randomly shuffled sequence, that should not be identified as phage by the tools. We have included the widely-used tool Kraken2 as an additional option for phage identification. The manuscript has been rewritten throughout to reflect the different outcomes with these datasets, compared to the data described in the previous version of our manuscript. We now provide guidance for users of tools to predict phage sequences in metagenomic datasets. Specifically we suggest that Kraken2 should be considered when researchers are trying to identify previously characterised phages. When novel phage detection is required Kraken2 should be used in combination with tools such as VirSorter2 and DeepVirFinder.

