## Supplementary Figure 1 for "Gauge your phage: Benchmarking of bacteriophage identification tools in metagenomic sequencing data"

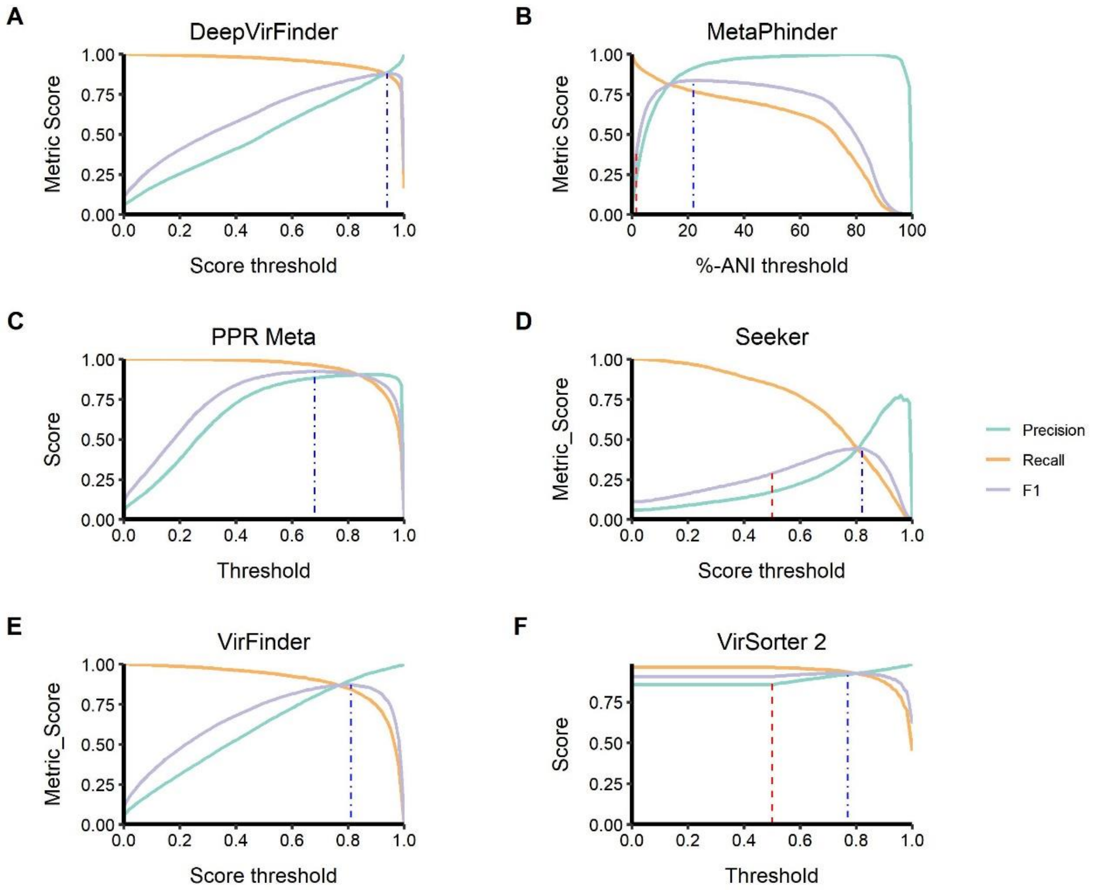


***Supplementary Figure 1: Tool optimisation curves***

*Optimisation curves for tools that provide score/probability thresholds. Precision, recall, and F1-score are plotted against threshold. Red dotted lines highlight the performance at a default threshold where provided by the authors, and blue dotted lines indicate the threshold that has the highest F1-score for each tool. A. DeepVirFinder. Contigs that are predicted as viral with a score below or equal to the threshold are used in the metric calculation. B. MetaPhinder. Contigs with an ANI-% greater or equal to the threshold are used in the metric calculation. C. PPR Meta. Contigs with a score higher or equal to the threshold are used to calculate the metrics. D. Seeker. Contigs with a score higher or equal to the threshold are used to calculate the metrics. E. VirFinder. Contigs that are predicted as viral with a score below or equal to the threshold are used in the metric calculation. F. VirSorter2. Contigs with a score higher or equal to the threshold are used to calculate the metrics.*
