## Supplementary Figure 2 for "Gauge your phage: Benchmarking of bacteriophage identification tools in metagenomic sequencing data"

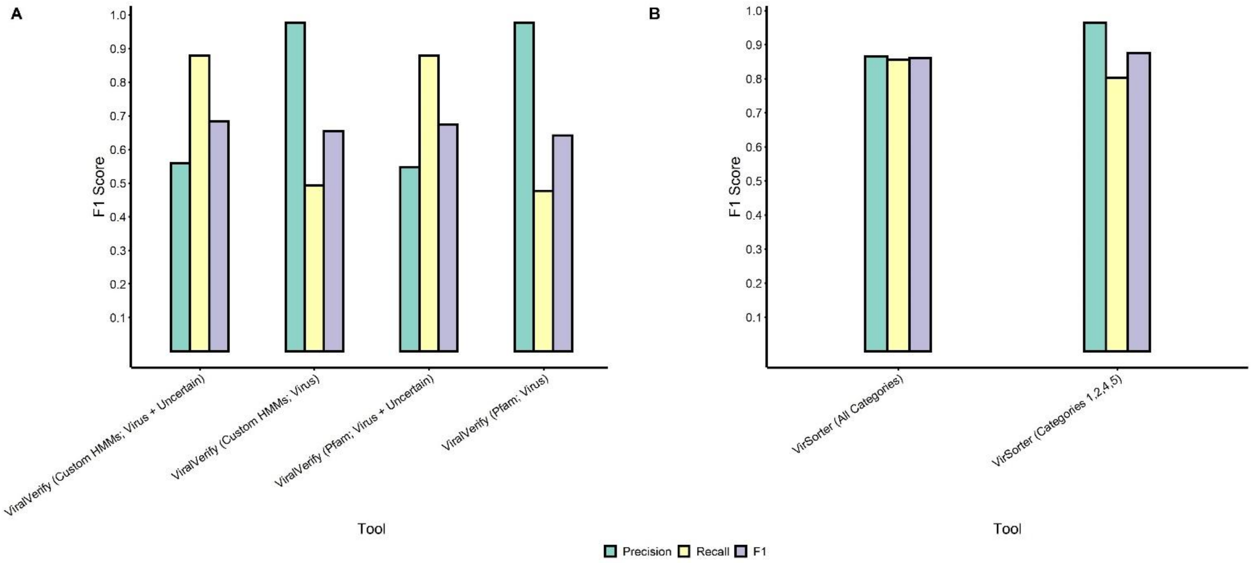


***Supplementary Figure 2: Tool category optimisations***

*F1-score plots of tools that provide categorical thresholds. A. viralVerify has two thresholds, Virus, and Virus Uncertain. This was also combined with two databases: Pfam 34.0 HMM database, and the custom database provided on viralVerify's GitHub repository. B. VirSorter was run with two category sets, the first included all six categories, the second taking the highest two categories from its viral prediction (category 1 and 2) and its prophage prediction (category 4 and 5).*
