## Supplementary Figure 3 for "Gauge your phage: Benchmarking of bacteriophage identification tools in metagenomic sequencing data"

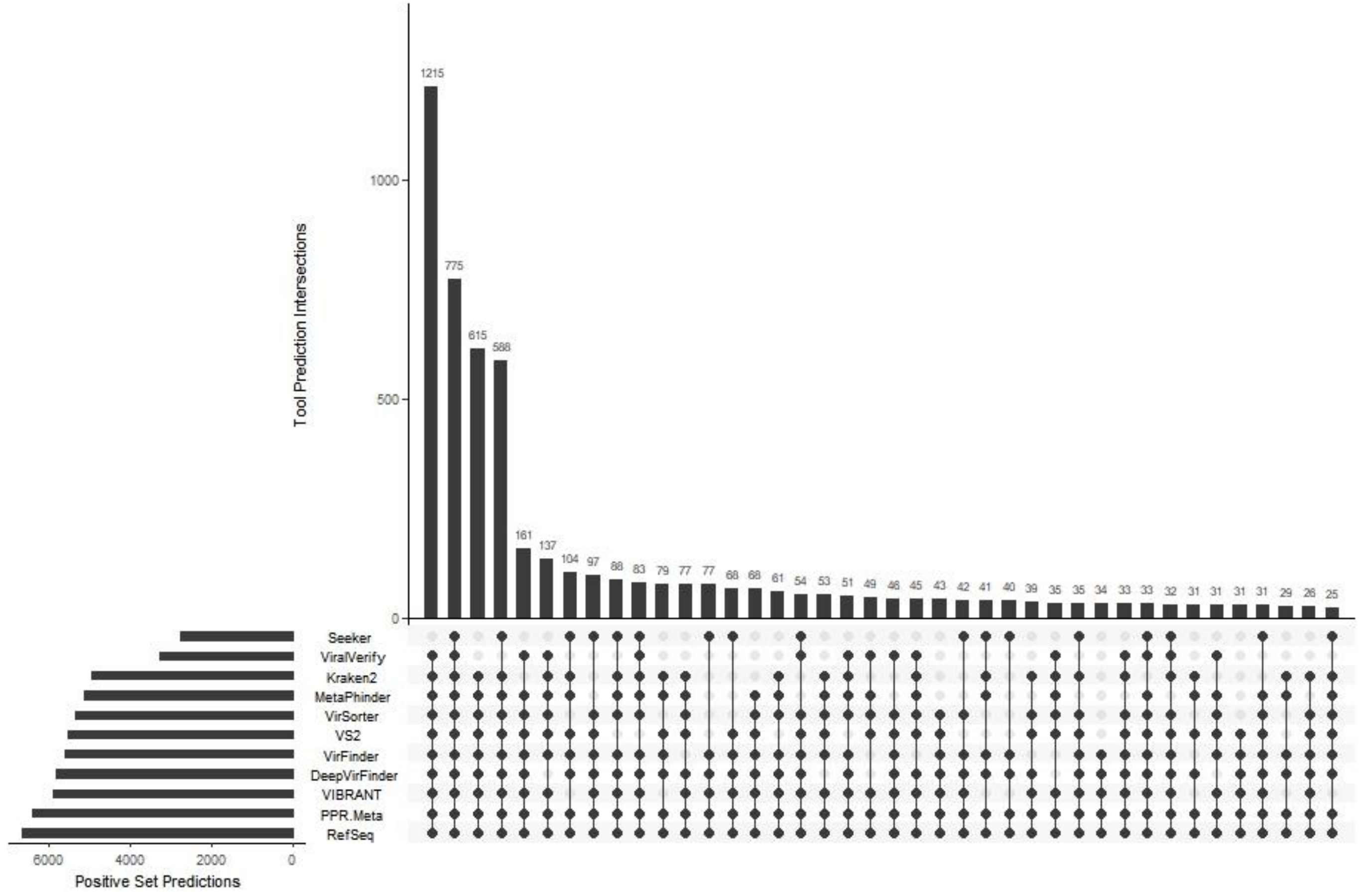


***Supplementary Figure 3: Upset plot of RefSeq phage artificial contigs predicted as viral by each tool.***

*The bar chart on the left indicates the total number of artificial RefSeq phage contigs predicted as viral by each tool. The upper bar chart shows the intersection size between contigs that were predicted by each tool, with the dark connected dots on the panel below indicating which tools were included in the intersection. The top 50 intersections are shown.*
