## Supplementary material for "Gauge your phage: Benchmarking of bacteriophage identification tools in metagenomic sequencing data": Table S1

***Supplementary Table 1: Bacteriophage strain composition of uneven mock community from Kleiner et al. [53]***

| Phage strain | Cell abundance (%) | Genome type | Host |
|---|---|---|---|
| ES18 | 0.363 | dsDNA | *Salmonella enterica* serotype Typhimurium LT2 |
| F0 | 2.925 | dsDNA | *Salmonella enterica* serotype Typhimurium LT2 |
| F2 | 0.250 | ssRNA | *Escherichia coli* K12 with Flac+ Plasmid |
| M13 | 0.250 | ssDNA | *Escherichia coli* K12 with Flac+ Plasmid |
| P22 | 21.250 | dsDNA | *Salmonella enterica* serotype Typhimurium LT2 |

*Composition, abundance, genome type and host of the five bacteriophage strains in the uneven mock community reported by (Kleiner et al., 2017).*
