## Supplementary material for "Gauge your phage: Benchmarking of bacteriophage identification tools in metagenomic sequencing data": Table S2

***Supplementary Table 2: Analysis of Similarity (ANOSIM) between non-metric multidimensional scaling of tools***

| **Tool** | **ANOSIM statistic R** | **P-value** |
|---|---|---|
| DeepVirFinder | 0.1264 | 0.1317 |
| Kraken2 | -0.07775 | 0.7357 |
| MetaPhinder | -0.0442 | 0.6041 |
| PPR Meta | -0.1073 | 0.8381 |
| **Seeker** | **0.3872** | **0.0028** |
| VIBRANT | -0.1028 | 0.8298 |
| VirFinder | 0.002688 | 0.4552 |
| VirSorter | -0.08822 | 0.7833 |
| VirSorter2 | -0.108 | 0.8375 |
| viralVerify | 0.02106 | 0.3985 |

*ANOSIM of each tool against all other tools and the default. Significance values reported are not corrected for multiple comparisons. Values deemed to be significant ($P < 0.05$) after adjusting the alpha value with the Benjamini–Hochberg method are highlighted in bold.*
